# Design of novel specific primers for helicobacter detection with diagnostic utility evaluation of different 16srRNA genus-specific primers: A comparative study

**DOI:** 10.1101/2021.11.29.470340

**Authors:** Shaymaa Abdelmalek, Karim Shokry, Wafy Hamed, Mohammed Abdelnalser, Ashraf Aboubakr, Sameh Abou Elenin, Mohamed Ali, Mohamed Mostafa, Mahmoud Abou-Okada

**Affiliations:** Microbiology department, Faculty of Veterinary Medicine, Cairo University; Microbiology department, Faculty of Veterinary Medicine, University of Sadat City; Internal Medicine and Gastroenterology, Military Medical Academy; Internal Medicine and Gastroenterology and Hepatology, Helwan Military production Hospital; Internal Medicine and Gastroenterology, Ahmed Maher Hospital, Cairo, Egypt; Aquatic Animal Medicine and Management, Faculty of Veterinary Medicine, Cairo University

**Keywords:** Helicobacter *spp.*, *H. pylori*, 16srRNA, ROC, PCR, nested, bioinformatics

## Abstract

Molecular diagnosis of helicobacters by PCR is simpler, more accurate, and feasible compared to other diagnostic methods. Validity and accuracy are highly dependent on the PCR primer design, diffusion time, and mutation rate of helicobacters. This study aimed to design 16srRNA -specific primers for Helicobacter *spp.* and *H. pylori*. Application of comparative statistical analysis of the diagnostic utility of the most available 16srRNA genus-specific primers. The new primers were designed using bioinformatics tools (MAFFT MSA and Gblocks command line). A comparative study was applied on nine genus-specific 16srRNA primers in comparison to the ConsH using Insilco and laboratory evaluation. The results demonstrated that the best specificity and sensitivity of the primers designed for this study compared to other primers. The comparative study revealed that the heminested outer/inner primers were the worst. Although H276,16srRNA(a), HeliS/Heli-nest, and Hcom had acceptable diagnostic utility, false positive and false negative results were obtained. Specificity testing on clinical samples indicated a surprising result; that *H. pylori* was not the sole enemy that we were looking for, but the NPH should be considered as a real risk prognostic for gastric diseases, consequently, a specific diagnosis and treatment should be developed. This study concluded that our designed primers were the most specific and sensitive in comparison with other primers. in addition, Insilco evaluation is not accurate enough for primer assessment and that the laboratory evaluation is mandatory.

## Introduction

Genus Helicobacter is a microaerophilic, helical, Gram-negative bacterium responsible for gastrointestinal disorders [1]. Thirty-four Helicobacter *spp.* have been identified so far according to LPSN (http://www.bacterio.net/h/helicobacter.html). It became evident that Helicobacter *spp.* can infect humans and various animal hosts and colonize different anatomical regions of the gastrointestinal tract [2].

*Helicobacter pylori* (*H. pylori*) was first recognized and considered a major cause of gastrointestinal disorders, including gastritis and peptic ulcers. Chronic infections are associated with gastric cancers and mucosal-associated lymphoid tissue lymphoma (type 1 gastric carcinogen by the International Agency for Cancer Research) in addition to several extra-gastric diseases [1, 3, 4].

Helicobacter *spp.* is assumed to be one of the most genetically diverse bacterial species studied to date [5]. Its genetic variability contributes to adaptation to changing environmental conditions via many approaches as antibiotic resistance development and the continuous surface antigen variation [6]. A wide range of genetic variability is attributed to substitution mutations [7], insertion elements [8], genetic rearrangement in the pathogenicity islands [9], the presence of prophages that causes genetic variations [10], and natural transformability demonstrated by many strains (11). Helicobacter *spp.* variability renders its diagnosis more challenging than many other genera.

Molecular diagnosis by polymerase chain reaction (PCR) is simpler, more accurate, and feasible compared to other invasive and non-invasive diagnostic procedures[12, 13, 14]. PCR is used for the detection of *H. pylori* DNA in various samples including gastric mucosa, feaces, saliva, dental plaque, and other environmental samples [15]. It has succeeded in demonstrating an actual correlation between *H. pylori* infection and extra gastric digestive carcinogenesis as hepatic carcinoma, bile duct cancer, pancreatic cancer, and colon cancer [16, 17].

Molecular diagnostic methods using PCR especially nested PCR will be the gold standard in helicobacter diagnosis [18]. Although their use is still mainly restricted to research, it is gaining great popularity in the medical field [19]. It is noteworthy that the validity and accuracy of results are highly dependent on the PCR design and the time of its publication as new sequences are constantly submitted to databases [16, 20].

Certain target genes have been extensively used for the PCR detection of Helicobacter *spp.* and *H. pylori*, including the 16S rRNA gene, the 26K species-specific antigen gene, the *glm*M gene, the *ure*A gene, the *ure*B gene, the *cag*A gene, and the *vac*A gene [21–33]. The most sensitive and widely used gene for the detection of Helicobacter infections is PCR that targets the genus-specific and conserved region of the housekeeping gene, 16S rRNA [16, 34], which is present in all bacterial species. This small ribosomal subunit gene contains conserved regions that are used for the general amplification of bacterial DNA by utilizing universal primers. Comparison of DNA sequences from these PCR products is widely used in taxonomy, phylogenetic studies (35), and clinical microbiology [36]. In addition to the conserved regions, 16S rRNA contains hypervariable regions that are highly specific for biological species or genera [37, 38].

Despite the various advantages of PCR, high mutation rates of Helicobacter *spp.*, the short primers, the low melting temperature, polymorphism in binding site at 3’end in protein-coding genes, and high melting temperature difference between forward and reverse primers (more than the recommended 4°C) affect amplification efficiency and may lead to false-negative results [18, 19, 39, 40]. False-positive results may be attributed to the usage of non-specific primers and the detection of cDNA from non-pyloric helicobacter strains (NPH), this is particularly important in environmental samples which may contain previously uncultured organisms or NPH [15, 41].

The present study aimed to design new sets of primers specific to Helicobacter *spp.* and *H. pylori* using bioinformatics tools. Evaluation of these designed primers was performed by Insilico and experimental testing. Comparative Statistical analyses were applied for most 16srRNA primers for Genus Helicobacter detection.

## Material and Methods

### 1. Design new 16srRNA specific primers for G: Helicobacter and *Helicobacter pylori*

#### Selected strains for multiple sequence alignment (MSA) and determination of the G: Helicobacter and *H. pylori* conserved regions

*Helicobacter pylori (2017, 2018, 26695, 26695, 35A, 51, 52, 83 908, 17, Aklavik86, B38, B8, BM0(12A, 12S), Cuz20, ELS37, F(16, 30, 32, 57), G27, Gambia94/24, HPAG1, HUP-B14, India7, Lithuania75 OK113 DNA, OK310, P12, PeCan (18, 4), Puno(120, 135), Rif(1, 2), SJM180, SNT49, Sat464, Shi(112, 169, 417, 470), South Africa(20, 7), UM(032, 037, 066, 298, 299), XZ274, v225d, strain J99.* Non-pyloric Helicobacter (NPH) *as Helicobacter acinonychis str. Sheeba, Helicobacter bizzozeronii CIII-1, Helicobacter cetorum (MIT 00-7128 and MIT 99-5656), Helicobacter cinaedi (ATCC BAA-847 DNA and PAGU611). Helicobacter felis ATCC 49179, Helicobacter hepaticus ATCC 51449, Helicobacter mustelae 12198*.

#### Multiple sequence alignment (MSA)

All 16srRNA gene sequences of all selected helicobacter strains were downloaded and saved in FASTA format to be used in MSA. MAFFT online version https://mafft.cbrc.jp/alignment/server/ was used for MSA of all 16srRNA gene sequences of the selected strains with a distance matrix by counting the number of shared 6mers between every sequence pair. A guide tree was built, and the sequences were aligned progressively according to the branching order, then the tree was reconstructed, and finally, a second progressive alignment was carried out. MAFFT was applied twice to obtain the outputs of Pearson/FASTA and Clustal.

#### Gblocks tool command line for the determination of conserved regions

To detect the conserved regions of the alignment, the Gblocks tool was used to yield an htm file that can be viewed using any browser. The MSA of all Helicobacter *spp.* 16srRNA gene sequences were collected in a file named **All_strains_16s.aln** and MSA of all *H. pylori* strains 16srRNA gene sequences were collected in a file named **All_H_pylori.aln. both files were used in The Gblocks command line in Terminal.**

$ Gblocks All_Strains_16s.aln -t=d -p=y

$ Gblocks All_H_pylori.aln -t=d -p=y

#### The design of new specific PCR primers for Helicobacter *spp.* (ConsH) and nested primers for *H. pylori* (PyloA/PyloAN)

The conserved regions of all Helicobacter *spp.* and *H. pylori* were used in Thermofisher oligonucleotides design online version https://www.thermofisher.com/eg/en/home/life-science/oligonucleotides-primers-probes-genes/custom-dna-oligos.html?s_kwcid=AL!3652!3!506722412345!p!!g!!thermo%20primer&ef_id=CjwKCAjw_o-HBhAsEiwANqYhp_8UC-ah9JbbfjH6_l3wIt2XPuxGgjhQjVGVJBB5ny9X8hWdECQ3hxoCa7oQAvD_BwE:G:s&s_kwcid=AL!3652!3!506722412345!p!!g!!thermo%20primer&cid=bid_mol_pch_r01_co_cp1358_pjt0000_bid00000_0se_gaw_bt_pur_con&gclid=CjwKCAjw_o-HBhAsEiwANqYhp_8UC-ah9JbbfjH6_l3wIt2XPuxGgjhQjVGVJBB5ny9X8hWdECQ3hxoCa7oQAvD_BwE, Which was used to design primers for PCR. Many probabilities were obtained for forward and reverse primers. Every pair was tested to select the most suitable pair.

### 2. Evaluation of the newly designed primers

#### Insilco evaluation by primer-Blast and Insilco PCR amplification

ConsH and PyloA/PyloAN primers were examined by primer-Blast online software https://www.ncbi.nlm.nih.gov/tools/primer-blast/. The primers were also subjected to Insilco PCR amplification using online software http://insilico.ehu.es/ against 62 Helicobacter strains including 53 *Helicobacter pylori* strains and 9 non-pyloric Helicobacter strains.

#### Laboratory evaluation

##### DNA extraction

The whole genome was extracted from three different samples, local isolate, gastric biopsies, and stool samples. DNA extraction using different extraction kits, Thermo scientific GeneJET Genomic DNA Purification kit#K0722, QIA*amp* DNA stool Mini kit#51604, and GF-1 Tissue DNA Extraction kit Vivantis cat.no:GF-TD-100

##### PCR

The PCR mixture included 1x master mix (amaR OnePCR^TM^, Cat.No. SM213-0250, GeneDireX, Inc.), 0.25 µmol forward and reverse primers, 10 ng DNA, and up to 20 µl nuclease-free water.

The amplification cycle for ConsH, pyloA/PyloAN was as follow: initial denaturation at 94°C/3 minutes, 30 cycles of the following temperatures: 94°C/30 sec, annealing at 60°C/1 minutes, and cyclic extension at 72°C/2 minutes, final extension at 72°C/5 minutes. Half a microliter of PCR product for the first nested PCR (PyloA) was amplified in the second PCR (PyloAN) for 20 cycles. Gel electrophoresis of the PCR products was performed in 1.5% agarose (Vivantis, cat.no: pco701) against a 100 bp DNA ladder (Vivantis, cat.no:NL1405).

##### Specificity testing

PCR application of newly designed primers using eight sequenced Helicobacter strains (5 *H. pylori* and 3 non-pyloric Helicobacter, *H. hepaticus*, *H. cinidiae,* and *H. felis*) and 10 non-Helicobacter bacteria present in GIT included *E. coli, Klebsiella pneumoniae, Salmonella spp., Proteus mirabilis, Enterococcus fecalis, Enterobacter spp., Staphylococcus aureus, Streptococcus mutans, Pseudomonas aeruginosa, and Candida albicans*. All bacterial strains were sequenced reference. The newly designed primers were applied clinically as mentioned before on 243 gastric biopsies taken from dyspeptic patients admitted to Ahmed Maher Hospital (180 samples), El-Maadi Military Hospital (38 samples), and Military production Hospital in Helwan (25 samples). The patients were tested in the hospital by different invasive methods like endoscopic pictures, histopathology, and rapid urea test (RUT). This study was approved by the Research Ethics Committee process number (HAM00116). The PCR positive samples were confirmed by the different invasive methods and sequencing of the PCR products.

#### Sensitivity testing

Serially diluted DNA of the whole genome of *H. pylori* (Local isolate) was introduced into a negative stool and biopsy samples in the following concentration (5 ng, 0.5 ng, 50 pg, 5 pg, 0.5pg/µl). The final concentration in total volume PCR mixture (250 pg, 25 pg, 2.5 pg, 0.25 pg, 25 fg) [42].

### 3. Comparative analysis between the newly designed primer ConsH and different 16srRNA primers for Helicobacter *spp.* detection

#### The 16srRNA specific primers used in the comparative study

The comparative study was performed on 9 primers for the detection of G: Helicobacter compared to the newly designed primer in this study (ConsH) (Table 1).

**Table 1:**
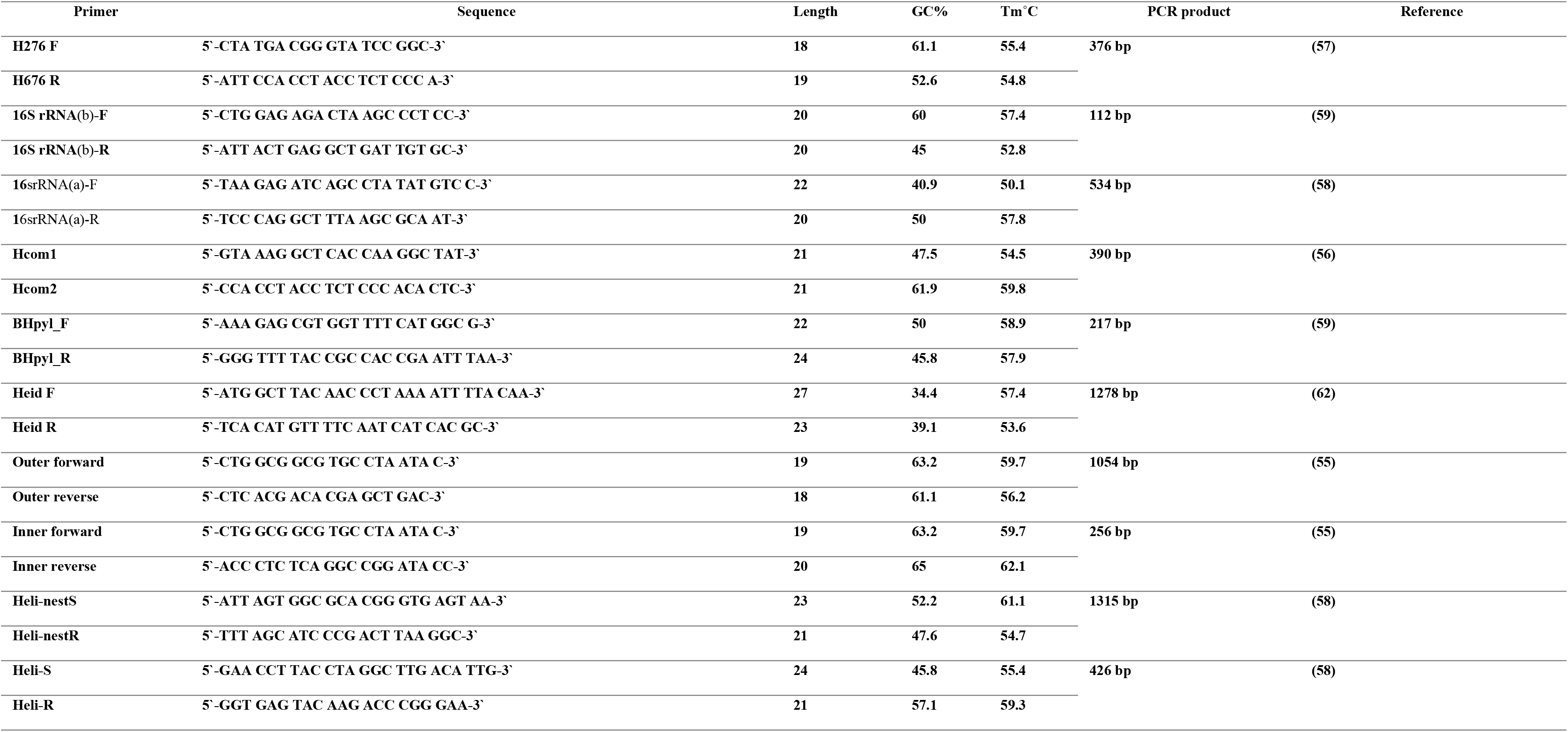
the sequence of different primers for detection of G: Helicobacter

#### Insilco comparative study

These primers were examined by primer-blast and subjected to Insilco PCR amplification.

#### Laboratory comparative study (specificity testing)

PCR application of the comparable primers was performed on the sequenced Helicobacter strains and non-helicobacter strains.

### 4. Statistical evaluation of the diagnostic utility of different 16srRNA primers for G: Helicobacter detection in the literature

#### Gold standard primer selection

The selected gold standard primer for evaluation was the most specific one which gave negative results with all non-Helicobacter strains and positive results with all sequenced Helicobacter strains.

##### Statistical methodology

The sensitivity (true positive rate, TPR), specificity (true negative rate, TNR), positive predictive value (PPV) and negative predictive value (NPV), false-positive rate (FPR), false-negative rate (FNR), positive likelihood ratio (LR+), negative likelihood ratio (LR−), accuracy (ACC), balanced accuracy (BA) and diagnostic odds ratio (DOR) were calculated for insilco and laboratory comparison of the nine comparable primers. Statistical analyses of screening tests for Helicobacter *spp*. using different primers were performed using Chi-square. Receiver Operating Characteristics (ROC) analysis for each screening test, the true positive rate (TPR) against false positive rate (FPR) can be measured compared to the gold standard. Values of *p* ≤ 0.05 were considered statistically significant. Analyses were performed via SPSS version 26 software (SPSS Inc, Chicago, IL, USA) and R version 4.1.1 (R Foundation for Statistical Computing), with the ‘epiR’ package [43, 44].

### Data availability

Helicobacter DNA sequences were submitted to NCBI Genbank with the following accession numbers OL630959, OL631133, OL631225, OL631585, OL634782, OL634838, OL631152, OL631161and, OL631249.

## Results

### 1. The newly designed 16srRNA specific primers for Helicobacter *spp.* and *H. pylori*

The conserved regions of the 16srRNA gene-specific to all helicobacters and all *H. pylori* strains are demonstrated in Table 2; whereas the conserved regions used in the design of highly specific primers are demonstrated in Table 3. The first conserved area of Helicobacter *spp.* was used for designing the genus-specific primer (ConsH) and the second conserved area of *H. pylori* was used for designing the *H. pylori* specific nested primer (PyloA/ PyloAN).

**Table 2:**
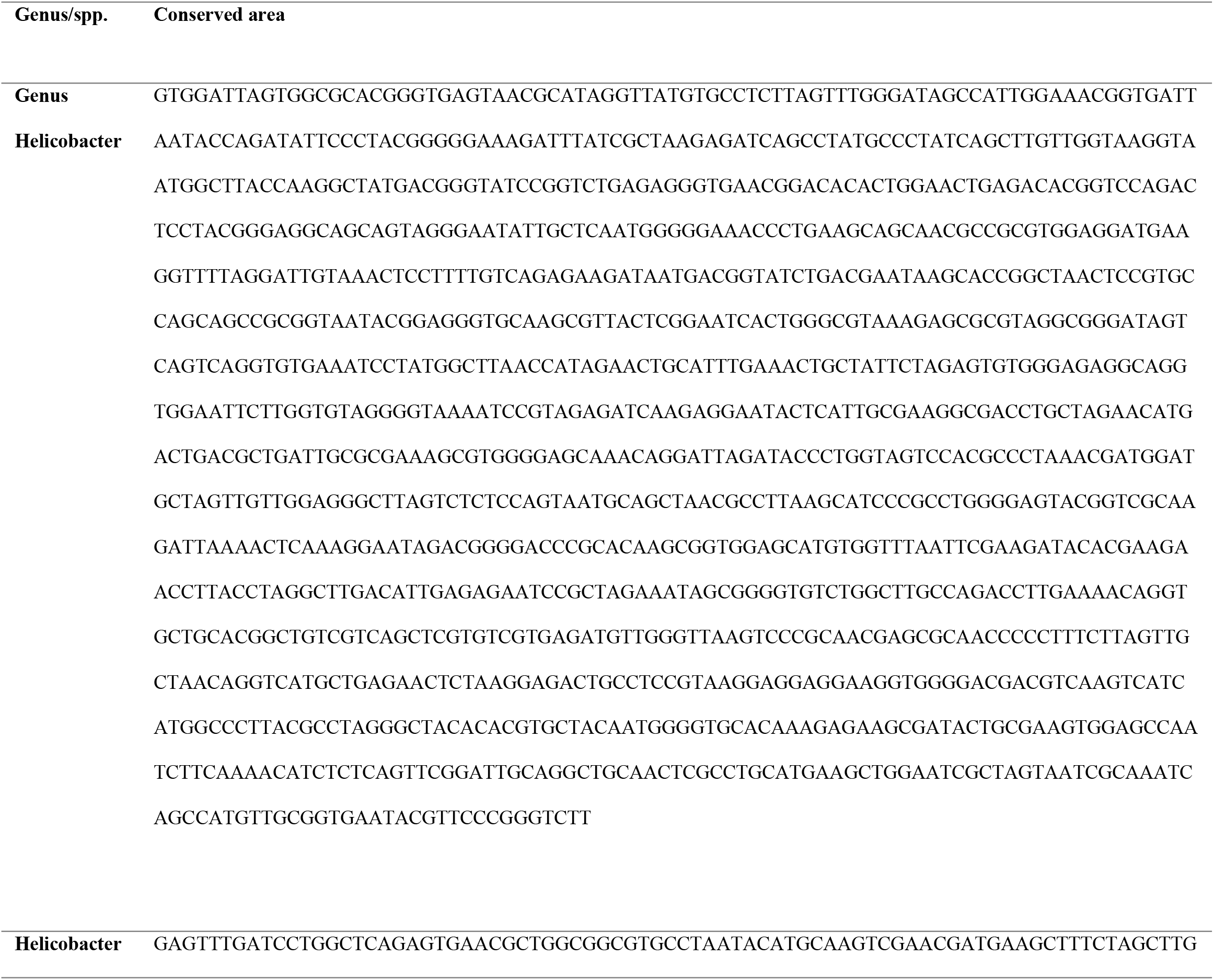
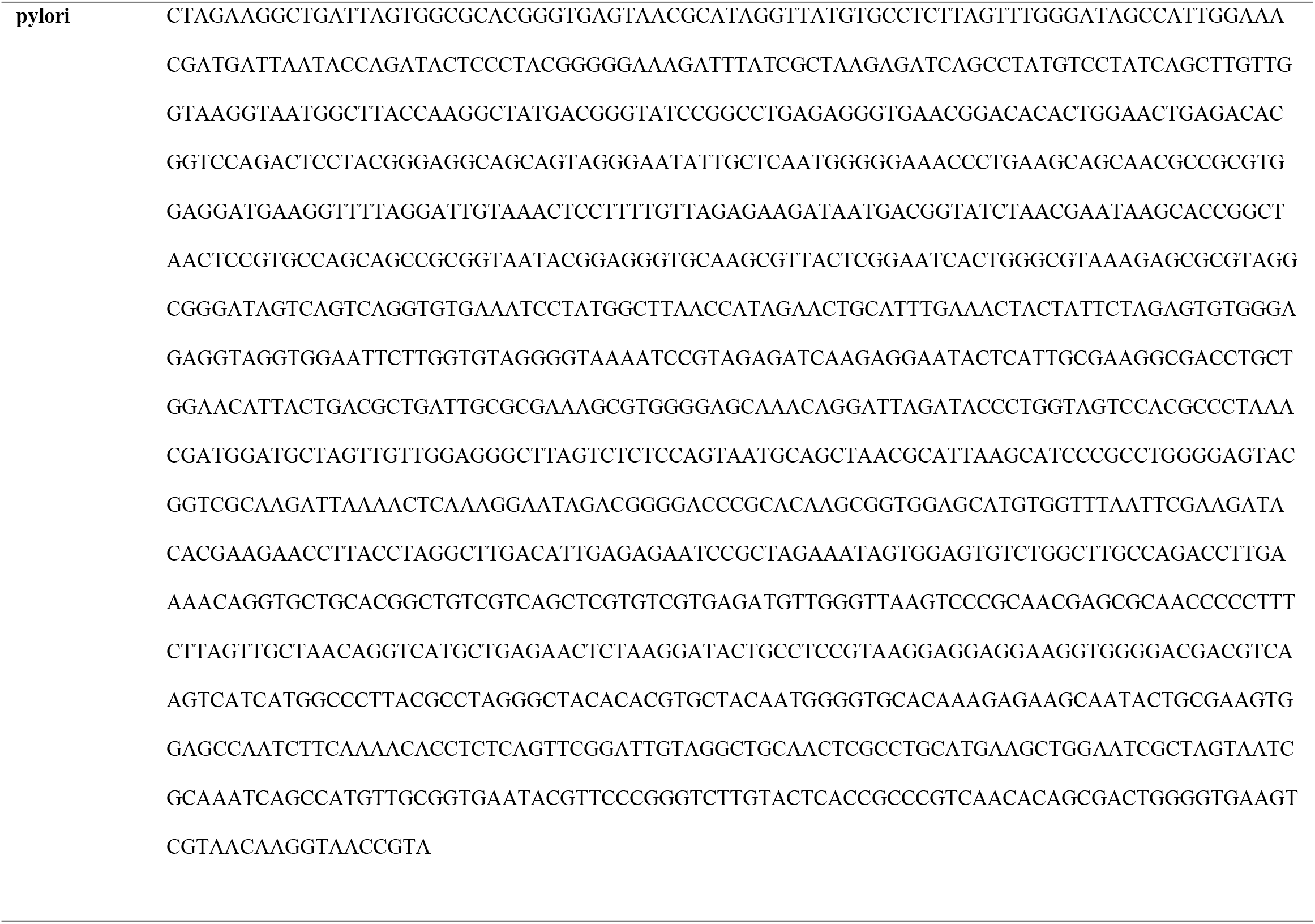
the conserved sequence of 16srRNA gene in G: Helicobacter and H. pylori

**Table 3:**
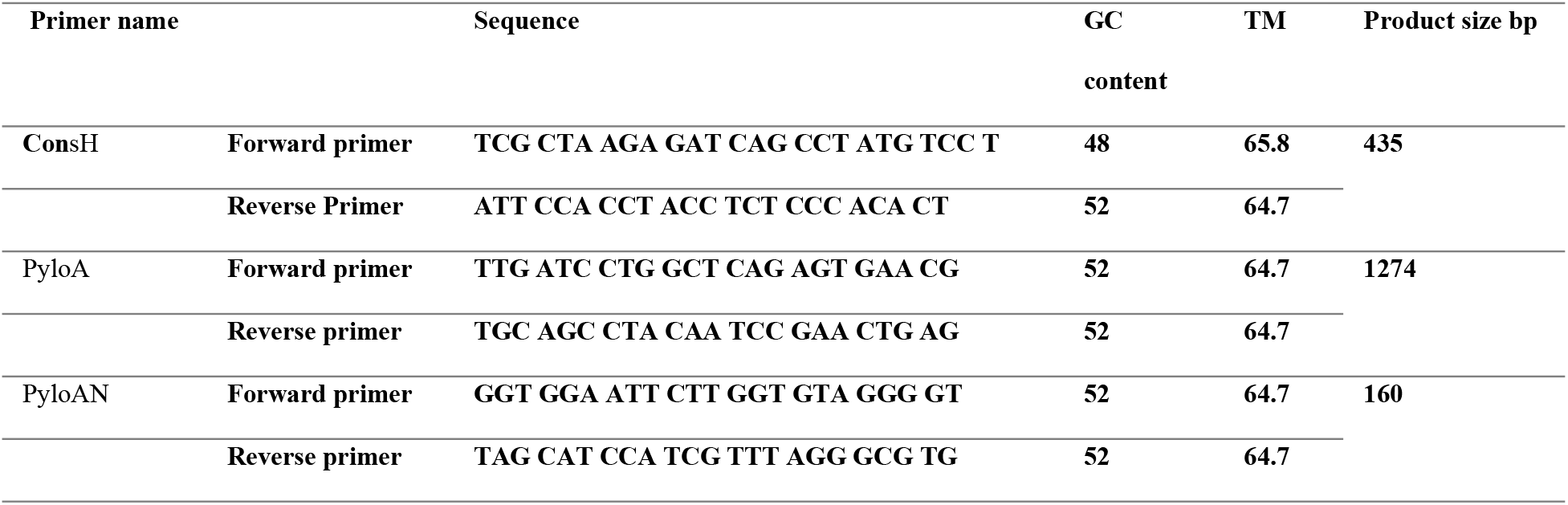
Sequence of the newly designed primers in the study

### 2. Evaluation of the newly designed primers

#### Insilco evaluation by BLAST and Insilco PCR amplification

ConsH primer corresponds to 383 *H. pylori* and 18 NPH by primer-blast analysis. The nested primer (PyloA/PyloAN) corresponds to 281 *H. pylori* strains. Insilco PCR amplification revealed that ConsH primer gave a positive band of 435 bp in 60 Helicobacter strains (96.8%). The nested primer (PyloA/PyloAN) revealed positive bands of 1274 bp (first nested, PyloA) and 160 bp (second nested, PyloAN) in 61 Helicobacter strains (98.4%).

#### Laboratory evaluation

- Specificity testing: The ConsH obtained positive bands of 435 bp with all helicobacters and negative results with all non-helicobacter strains. The H. pylori-specific nested primer (PyloA/PyloAN) produced positive bands of 1274 bp in the first pair and 160 bp in the second pair with all five H. pylori strains and negative results with all NPH and non-helicobacter strains. The results of PCR application in 243 clinical biopsy samples revealed 99 positive samples for Helicobacter spp. by ConsH primer (40.7%), from these positive samples, 66% were H. pylori and 33% were NPH. The positive cases were confirmed by sequencing (Fig 1).
- Sensitivity testing: The detection limit of ConsH primer was 250 pg and 25pg in stool and biopsy clinical samples, respectively, while the detection limit of PyloA was 0.25 fg and 0.25pg in stool and biopsy clinical samples respectively (Fig 2).

### 3. Comparative analysis between the newly designed primer ConsH and different 16srRNA primers for G: Helicobacter detection

#### Insilco comparative analysis

Primer Blast revealed that H276 primer correlates with 277 *H. pylori* and 56 NPH, whereas 16srRNA(b) matches 401 *H. pylori*, outer/inner primers are compatible with 283 *H. pylori* and 48 NPH, HeliS-Helinest match 329 *H. pylori*, and 115 NPH. 16srRNA(a) doesn’t match any Helicobacter. Hcom corresponds to 282 *H. pylori* and 13 NPH. BFHpyl matches 312 *H. pylori* and 3 NPH. Heid correlates with 297 *H. pylori* and 2 NPH. The positive percent of Insilco PCR amplification is explained as follows: H276 and outer/inner primers gave positive in 60 Helicobacters (96.8%), Hcom1 gave positive with 61 Helicobacters (98.4%). 16srRNA (a) and BFHpyl gave positive with only one Helicobacter (*H. pylori* 26695 and *H. pylori* G27, respectively). 16srRNA (b) and Heid didn’t produce any bands with all Helicobacters.

**Figure 1.**
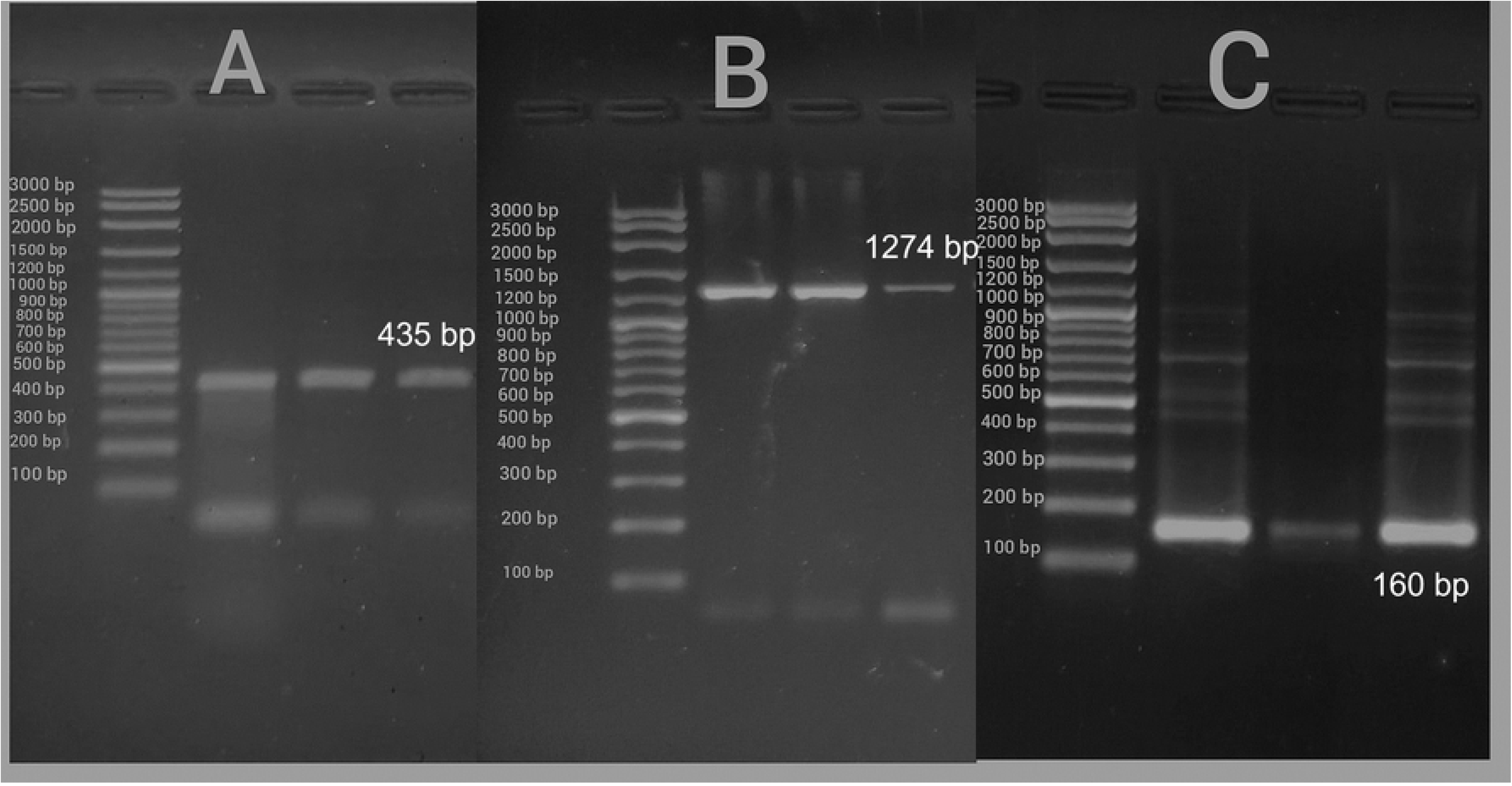
Gel electrophoresis of PCR products of A. ConsH amplified products(435bp) B. PyloA amplified products (1274bp) C. PyloAN amplified products (160 bp) in reference to 100 bp DNA ladder.

**Figure 2.**
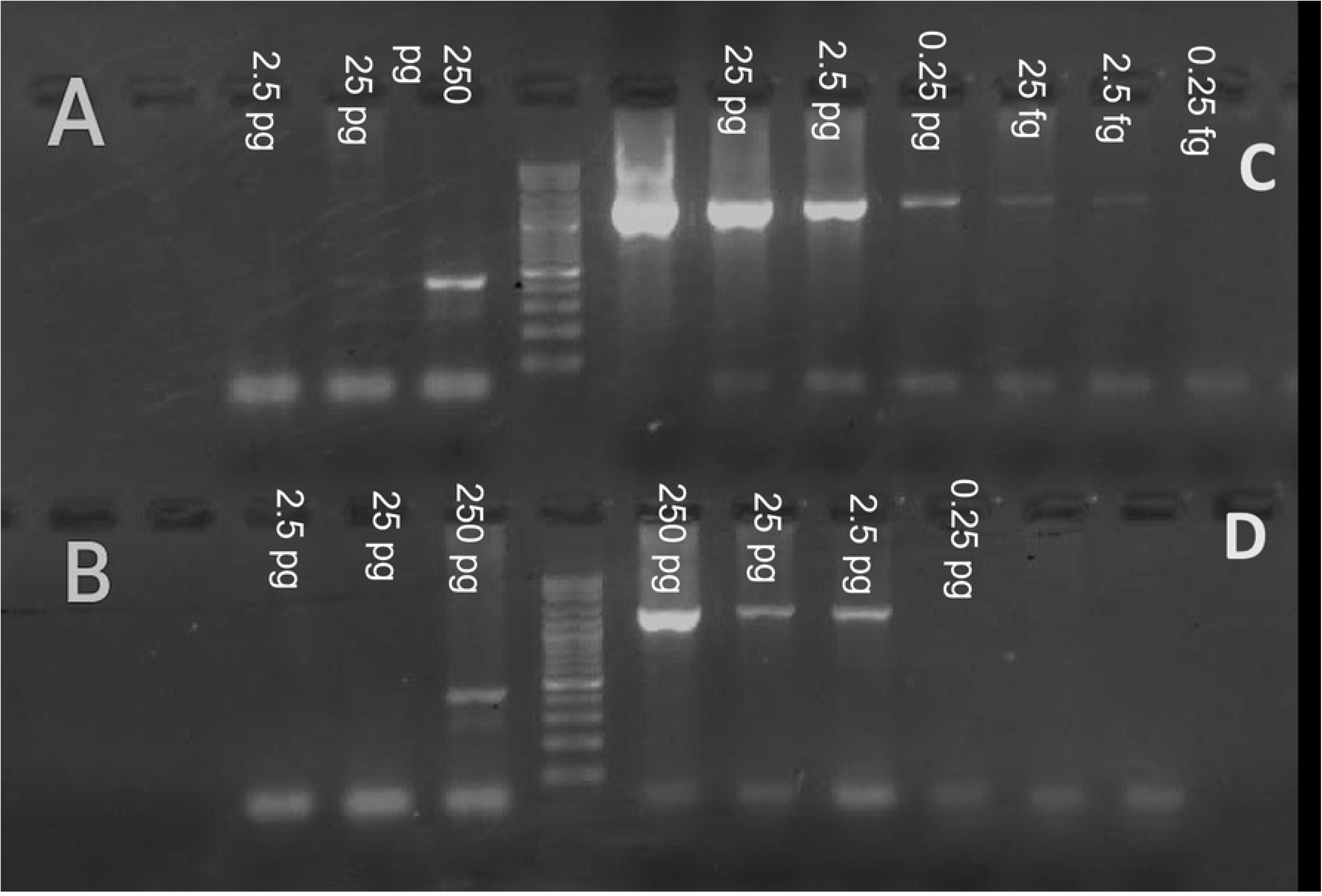
A, Sensitivity testing of primers ConsH and C, Sensitivity testing of PyloA using Stool clinical sample DNA with different concentrations. B, Sensitivity testing of Primers ConsH and D, Sensitivity testing of PyloA using biopsy clinical sample DNA with different concentrations.

#### Laboratory comparative study

H276, outer/inner, Hcom1, Heli-nest/HeliS, and 16srRNA (a) yielded positive results with all eight Helicobacters, 16srRNA (b) relented positive results with six Helicobacters, BFHpyl revealed a positive band with only one *H. pylori* while Heid primer gave negative with all Helicobacter *spp.* Nonspecific results were obtained with other Non-helicobacters as follows: H276 (*S. mutans)*, Heli-nest/HeliS (*Salmonella spp.*), Hcom1 and BFHpyl (*K. pneumonae)*, 16srRNA (a) (*E. fecalis*), Heid (*E. coli)* while outer/inner primer gave positive results with all non-helicobacters.

#### Statistical evaluation of the diagnostic utility of different 16srRNA primers for Helicobacter *spp.* detection

ConsH matched all criteria considered for the gold standard selection. The statistical evaluation parameters according to Insilco PCR amplification are described in Table 4. The outer/inner primers had 100% specificity and sensitivity with a *P*-value of 0.0001. Hcom1, H276, and HeliS/Heli-nest had 50% SP. The utility is measured also by the receiver operating characteristics (ROC) analysis expressed by the area under the curve (AUC), the ROC analysis of the Insilco evaluation is demonstrated in Fig 3. The laboratory evaluation showed 0.00% SP with heminested outer/inner primers with *P*-value=1. The H276, 16srRNA(a), HeliS/Heli-nest, and Hcom had 90% specificity with *P*-value= 0.001. Although 16srRNA(b) had 100% specificity, it had 75% sensitivity with a *P*-value=0.004. The ROC analysis of laboratory comparison is demonstrated in Fig 4.

**Figure 3.**
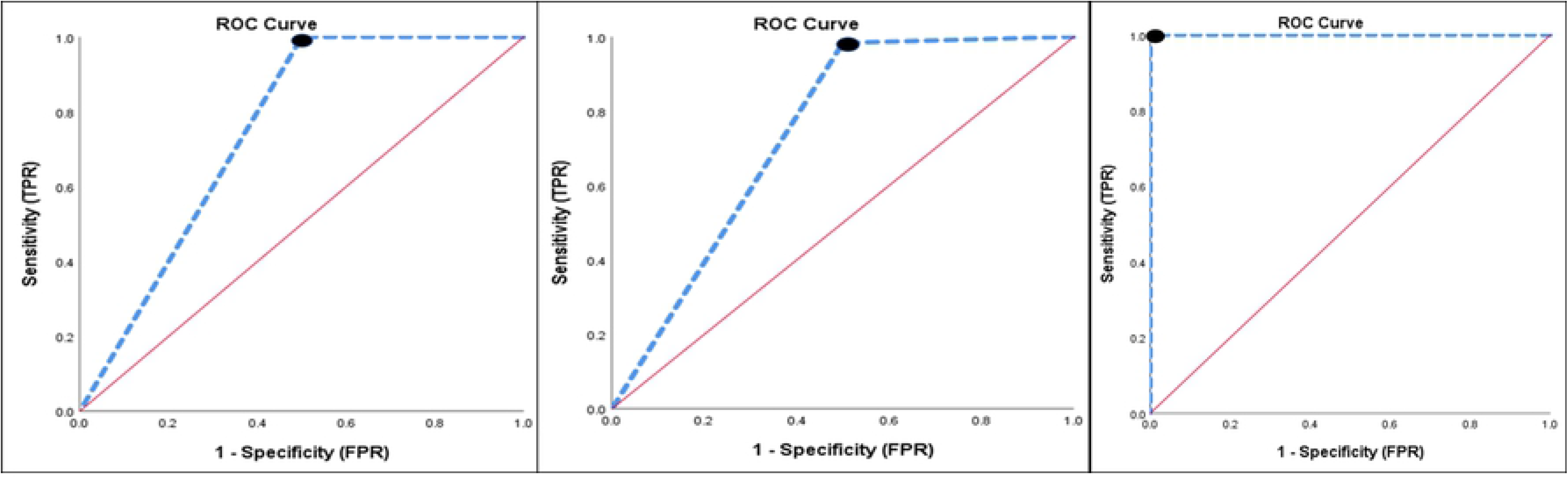
ROC curve of Insilco comparison by PCR amplification. Null hypothesis: true area = 0.5, AUC: A, ROC curve of ConsH and Hcom1 primers for detection of 62 Helicobacter strains, AUC represents the accuracy of the screening test (0.750). B, ROC curve of ConsH and H276 primers, AUC represents the accuracy of the screening test (0.742).C, ROC curve of ConsH and Outer + Inner primers, AUC represents the accuracy of the screening test (1.00).

**Figure 4.**
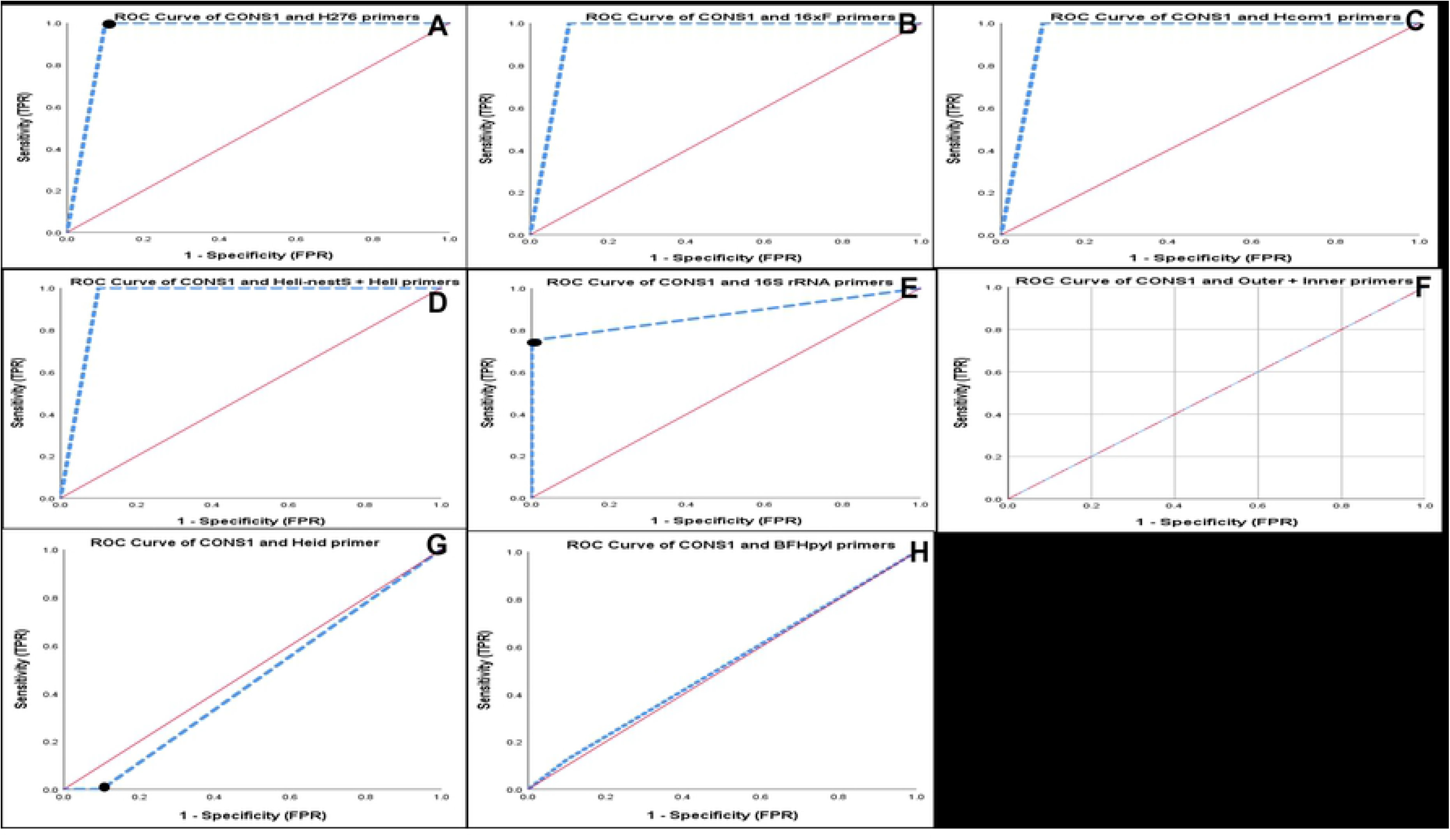
ROC curve of laboratory comparison of all primers using Helicobacter spp. (n= 8) and non-Helicobacter spp. (n= 10). Null hypothesis: true area = 0.5, AUC: Area under the curve. A, ROC curve of ConsH and H276 primers, AUC represents the accuracy of the screening test (0.950). B, ROC curve of ConsH and 16srRNA (a) primers, AUC represents the accuracy of the screening test (0.950). C, ROC curve of ConsH and Hcom1 primers, AUC represents the accuracy of the screening test (0.950). D, ROC curve of ConsH and Heli-nestS + Heli primers, AUC represents the accuracy of the screening test (0.950). E, **ROC curve of ConsH and 16S rRNA primers**, AUC represents the accuracy of the screening test (0.875). F, **ROC curve of ConsH and Outer + Inner primers**, AUC represents the accuracy of the screening test (0.500). G, **ROC curve of ConsH and Heid primers**, AUC represents the accuracy of the screening test (0.450). H, **ROC curve of ConsH and BFHpyl primers**, AUC represents the accuracy of the screening test (0.513).

**Table 4:** Screening primers test results of insilico PCR for detection of 62 Helicobacter strains.

| Gold<br>standard<br>Primer | Screening<br>Primers | SEN<br>(%) | SPC<br>(%) | PPV<br>(%) | NPV<br>(%) | FPR<br>(%) | FNR<br>(%) | Acc<br>(%) | BA<br>(%) | LR+ | LR- | DOR | X<sup>2</sup> | P value |
| --- | --- | --- | --- | --- | --- | --- | --- | --- | --- | --- | --- | --- | --- | --- |
| CONSH | Hcom1 | 100 | 50 | 98.4 | 100 | 50 | 0.0 | 98.4 | 75 | 2.0 | 0.0 | - | 7.12 | 0.01 |
|  | H276 | 98.3 | 50 | 98.3 | 50.0 | 50.0 | 1.7 | 96.8 | 74.2 | 1.97 | 0.033 | 59 | 3.140 | 0.076 |
|  | Outer + Inner | 100 | 100 | 100 | 100 | 0.0 | 0.0 | 100 | 100 | NA | 0.0 | - | 34.10 | 0.0001 |
|  | Heli-nestS +<br>Heli | 8.3 | 50 | 83.3 | 1.8 | 50 | 91.7 | 9.7 | 29.2 | 0.17 | 1.83 | 0.09 | 0.555 | 0.456 |

SEN: sensitivity; SPC: specificity; PPV: Positive predictive value; NPN: Negative predictive value; FPR: False positive rate; FNR: False-negative rate; ACC: Accuracy; BA: Balanced accuracy; LR+: Positive likelihood ratio; LR-: Negative likelihood ratio; DOR: Diagnostic odds ratio. 16S rRNA, hspG, and Heid primers tested negative in the detection of all 62 *Helicobacter* species. 16S rRNA and BFHpyl primers tested negative in the detection of 61 *Helicobacter* species.

## Discussion

The present study demonstrated the whole process of designing novel sets of highly specific primers targeting 16srRNA conserved region by bioinformatics tools, ConsH for detecting G: Helicobacter, and nested primers for *H. pylori* detection (PyloA/PyloAN). The design process was followed by evaluation of the diagnostic utility of the primers in comparison to a group of widely used 16srRNA primers in literature, and statistical monitoring of the diagnostic utility of the comparable primers used in Helicobacter *spp.* diagnosis.

Efficient PCR performance is highly dependent on PCR primer design, consequently one must spend a notable effort on the primer design. Well-designed PCR primers not only augment specificity and sensitivity but also reduce the effort spent on the experimental optimization [45].

PCR diagnosis especially nested PCR may be regarded as the gold standard for Helicobacter diagnosis through the construction of specific primers. Nested PCR provides higher sensitivity by excluding false-negative results due to low bacterial counts and PCR inhibitors [18]. PCR yielded a higher detection rate (40.8%) compared to histopathology (36.7%) and can be suitable for patients unfit for endoscopic examination [46].

The choice of 16sRNA gene in PCR diagnosis of Helicobacter diseases was supported by a systematic review and meta-analysis conducted on various sources, including MEDLINE, Web of Sciences, and the Cochrane Library from April 1, 1999, to May 1, 2016. The most diagnostic candidate genes according to statistical parameters were 23S rRNA, 16S rRNA, and *glm*M [47]. The urgent demand for rapid accurate Helicobacter detection puts an obligation for the construction of novel specific primers.

Both the ConsH and PyloA/PyloAN primers were synthesized according to the bioinformatic tools with primer designs criteria which ensures efficient PCR performance. The primer’ lengths fall within the recommended 18-30 nucleotides, shorter primers can produce non-specific results, and longer primers can form secondary structures and reduce PCR efficiency. Primers GC contents are within 48-52 %, and the difference between the melting temperatures of the forward and reverse primers was within 4℃ versus other compared primers as 16srRNA (a), Hcom, and Heli-nest primers which have larger than 4°C difference between the primers. This may cause false negatives [48, 49].

The evaluation of the newly designed primers was performed by Insilco tools and laboratory test (PCR). The results of the Insilco evaluation were very promising as ConsH matched with Helicobacters only and no mismatching with other bacteria, also the nested primers for *H. pylori* (PyloA/PyloAN) matches only with *H. pylori*. Insilco PCR amplification showed highly encouraging results. The advantages of Insilco evaluation are low cost and timesaving for evaluation as it gives a preliminary decision about the newly designed primer sets. Nonetheless, the Insilco evaluation is not a confirmatory method for evaluation due to continuously submitting uncurated sequences into GenBank, therefore, laboratory evaluation is mandatory [50].

In the laboratory evaluation, the newly designed primers showed sufficient results. The specificity testing revealed significant results as ConsH detected all Helicobacters and the nested primer (PyloA/PyloAN) identified all five *H. pylori* strains without mismatching with Non-pyloric Helicobacters. They did not select any non-Helicobacter bacteria. The clinical precision testing by PCR was applied on gastric biopsies from dyspeptic patients introduced to the endoscope unit in the mentioned hospitals, revealing that all PCR results are consistent with the invasive methods applied (RUT and Histopathology). The positive PCR by sequencing confirmed the presence of Helicobacter DNA in the samples [51–53]. The ConsH detected 40.7% positive samples, from the positive Helicobacter *spp.* PCR, the nested PCR detected 61.3% *H. pylori* and 39.7% NPH. These findings are concerning because there is an unexpected hidden unexpected enemy in the form of NPH, hence, diagnosis and treatment guidelines should be changed to consider the NPH beside *H. pylori*.

In the present study, the designed primers showed considerable sensitivity to low concentrations of specific DNA in clinical samples, allowing sensitive detection in different types of contaminated samples [54].

Here comes the answer to an important question about the need for a newly designed set of specific primers instead of the primers used in the literature. Insilco and laboratory comparative analysis should be established to determine the diagnostic utility of 16srRNA primers for genus-level identification of Helicobacter *spp*. Insilco comparison showed reasonable results for H276, 16srRNA(a), heminested outer/inner, and nested HeliS-Helinest by primer-blast but did not achieve substantial findings with 16srRNA(b). Insilco PCR amplification gave promising results with H276(96.8%), Hcom (98.4%), nonetheless, the results were poor for 16srRNA(b), BFHpyl, 16srRNA(a), and Heid primers. From the mentioned results, the Insilco evaluation is not an accurate method for a real evaluation of any primer but is only a preliminary step, so laboratory evaluation is the confirmatory method for accurate evaluation. The specificity test revealed that ConsH primer is the best one in this marathon, as it has all considerations to be the gold standard primer for statistical evaluation of the comparable primers. The results of the laboratory concluded that the comparable primers offered false-positive results by non-specific binding to non-Helicobacter bacteria and some of them gave false-negative results as BFHpyl, Heid, and 16srRNA(b).

The comparative study was analyzed statistically to assess the diagnostic utility. The diagnostic parameters and ROC analysis of Insilco and laboratory evaluation are demonstrated in Tables 4, 5, and Figs 3, 4, which concluded that the Insilco evaluation is not accurate enough to assess the diagnostic utility of any primer. That was evident in the case of heminested outer/inner primer which showed excellent diagnostic significance (*P-* 0.0001), 100% in most diagnostic parameters, and the best score of AUC (1.00) in ROC analysis. These results differ in the laboratory evaluation. This heminested primer produced false-positive results because it achieved positive PCR with all non-Helicobacter bacteria with 0.00% specificity (Table 5) and AUC (0.5) (Fig 4), which demonstrated the unreliability of heminested outer/inner primer for G: Helicobacter diagnosis. The aforementioned findings disagree with Qin *et.al*., who confirmed its reliability for G: Helicobacter identification and that it is a powerful diagnostic tool [55]. Hcom1, H276, and nested primer (HeliS/Heli-nest) indicated non-significant results for their use in G: Helicobacter diagnosis by Insilco PCR amplification (Table 4) and considered insignificant tool for diagnosis, as they had AUC less than 0.8 in ROC analysis tool. The laboratory evaluation of these primers demonstrated a major difference with Insilco evaluation as the statistical analysis revealed that they had significant diagnostic utility (*P-*0.001) and were considered reliable for Helicobacter *spp.* diagnosis with AUC (0.950) in ROC analysis [56–58]. However, the results of the current study are incompatible with Flahou *et al.,* that Hcom1 is suitable for the identification of Helicobacter *spp.* [59], as it may give false-positive results with *K. pneumonea*. H276 may give positive results with *S. mutans* found in the buccal cavity [60]. The present study disagrees with Riley *et al.,* who developed this primer and concluded that it was sensitive and specific to detect several numbers of Helicobacters and it was suitable for routine diagnosis [57]. The nested primers (HeliS/Heli-nest) gave positive results with Salmonella *spp.* which is alarming due to misdiagnosis of gastrointestinal disturbance of Helicobacteriosis and Salmonellosis [58].

**Table 5:** Screening primers results for detection of Helicobacter spp. (n= 8) and Non-Helicobacter spp. (n= 10)

| Gold<br>standard<br>Primer | Screening<br>Primer | SEN<br>(%) | SPC<br>(%) | PPV<br>(%) | NPV<br>(%) | FPR<br>(%) | FNR<br>(%) | Acc<br>(%) | BA<br>(%) | LR+ | LR- | DOR | X<sup>2</sup> | P value |
| --- | --- | --- | --- | --- | --- | --- | --- | --- | --- | --- | --- | --- | --- | --- |
| CONSH | H276 | 100 | 90.0 | 88.9 | 100.0 | 10.0 | 0.0 | 94.4 | 95 | 10.0 | 0.0 | - | 11.025 | 0.001 |
|  | 16 XF | 100 | 90.0 | 88.9 | 100.0 | 10.0 | 0.0 | 94.4 | 95 | 10.0 | 0.0 | - | 11.025 | 0.001 |
|  | Heli-nestS +<br>Heli | 100 | 90.0 | 88.9 | 100.0 | 10.0 | 0.0 | 94.4 | 95 | 10.0 | 0.0 | - | 11.025 | 0.001 |
|  | Hcom1 | 100 | 90.0 | 88.9 | 100.0 | 10.0 | 0.0 | 94.4 | 95 | 10.0 | 0.0 | - | 11.025 | 0.001 |
|  | 16S rRNA | 75.0 | 100 | 100 | 83.3 | 0.0 | 25 | 88.9 | 87.5 | - | 0.25 | - | 8.128 | 0.004 |
|  | Outer + Inner | 100 | 0.0 | 44.4 | - | 100 | 0.0 | 44.4 | 50 | 1.0 | - | - | - | - |
|  | BFHpyl | 12.5 | 90 | 50 | 56.3 | 10 | 87.5 | 55.6 | 51.3 | 1.25 | 0.97 | 1.29 | 0.00 | 1 |
|  | Heid | 0.0 | 90 | 0.0 | 52.9 | 10 | 100 | 50.0 | 45 | 0.0 | 1.11 | 0.0 | 0.00 | 1 |

SEN: sensitivity; SPC: specificity; PPV: Positive predictive value; NPN: Negative predictive value; FPR: False positive rate; FNR: False-negative rate; ACC: Accuracy; BA: Balanced accuracy; LR+: Positive likelihood ratio; LR-: Negative likelihood ratio; DOR: Diagnostic odds ratio.

16srRNA (a,b) were unreliable based on Insilco evaluation, but have a significance for diagnosis of Helicobacters (*P-* 0.001 and 0.004, respectively) and an acceptable AUC (0.875, 0.950, respectively) in ROC analysis. However, false-positive results may be detected in 16srRNA (b) with *E. fecalis.* Low sensitivity obtained with 16srRNA(a) (Se 75%). Consequently, these results disagree with Idowu *et al.,* and tiwari *et al.,* who concluded their specificity and sensitivity in Helicobacter *spp.* diagnosis [58, 61]. Finally, the worst primers that appeared in our study were clearly demonstrated in BFHpyl and Heid which have the lowest significance in Insilco and laboratory evaluation (*P-*1.00) and low AUC (0.513, 0.450, respectively) by ROC analysis (Table 5) (Fig 3, 4). These results disagree with Flahou *et al.,* and Farshad [59, 62].

Our study explained a suitable comparative analysis of different primers for Helicobacters diagnosis by using ROC analysis and different statistical diagnostic parameters. Use of bioinformatics tools and command line to extract conserved regions of 16srRNA gene in Helicobacter *spp.* and *H. pylori* are used in the design of primers. Considerable detailed evaluation of the newly designed primers by specificity using sequenced Helicobacters and non-Helicobacters carefully selected from GIT microbes. Application of the designed primers in clinical samples (gastric biopsies) using sufficient representative sample size from dyspeptic patients introduced to three hospitals. The results were compared with the routine diagnosis in the hospitals and sequencing of the PCR products. Sensitivity testing was performed to find the detectable DNA concentration in highly contaminated samples (Stool). However, more non-Helicobacters and sequenced local Helicobacter strains should be used, and that will be considered in our future studies.

## Conclusion

Our designed primers ConsH and PyloA/PyloAN are highly sensitive and specific and can be used for accurate diagnosis of Helicobacter diseases from different clinical samples. Moreover, PyloA/PyloAN indicated the abundance of non-pyloric Helicobacters and their unrecognized role in Helicobacteriosis. It can also be concluded that Insilco evaluation is not adequately accurate to assess the diagnostic utility of the primers and must be accompanied by the laboratory evaluation.

## Acknowledgment

This study was supported by the hospitals which supported us with the clinical samples, Ahmed Maher Hospital, Millitary production hospital in Helwan, and El-Maadi Millitary hospital, Cairo, Egypt.

## Notes

### Competing Interest Statement

The authors have declared no competing interest.

